# Morphine exposure during adolescence induces enduring social changes dependent on adolescent stage of exposure, sex, and social test

**DOI:** 10.1101/2023.04.21.537856

**Authors:** David N. King’uyu, Erin L. Edgar, Christopher Figueroa, J.M. Kirkland, Ashley M. Kopec

## Abstract

Drug exposure during adolescence, when the ‘reward’ circuitry of the brain is developing, can permanently impact reward-related behavior. Epidemiological studies show that opioid treatment during adolescence, such as pain management for a dental procedure or surgery, increases the incidence of psychiatric illness including substance use disorders. Moreover, the opioid epidemic currently in the United States is affecting younger individuals raising the impetus to understand the pathogenesis of the negative effects of opioids. One reward-related behavior that develops during adolescence is social behavior. We previously demonstrated that social development occurs in rats during sex-specific adolescent periods: early to mid-adolescence in males (postnatal day (P)30-40) and pre-early adolescence in females (P20-30). We thus hypothesized that morphine exposure during the female critical period would result in adult sociability deficits in females, but not males, and morphine administered during the male critical period would result in adult sociability deficits in males, but not females. We found that morphine exposure during the female critical period primarily resulted in deficits in sociability in females, while morphine exposure during the male critical period primarily resulted in deficits in sociability primarily in males. However, depending on the test performed and the social parameter measured, social alterations could be found in both sexes that received morphine exposure at either adolescent stage. These data indicate that when drug exposure occurs during adolescence, and how the endpoint data are measured, will play a large role in determining the effects of drug exposures on social development.

## 1. INTRODUCTION

Drug exposure during adolescence, when the ‘reward’ circuitry of the brain is developing, can permanently impact reward-related behavior into adulthood [1–8]. Morphine is an opioid primarily used for pain management by producing analgesic effects and is clinically indicated when non-narcotic pain management cannot be achieved [9–12]. Epidemiological studies show that opioid treatment during adolescence, such as pain management for a dental procedure or surgery, increases the incidence of psychiatric illness [13–15] including substance use disorders [16–18] in some reports by as high as 30% [19]. Moreover, the opioid epidemic currently in the United States is affecting younger individuals [20, 21] raising the impetus to understand the pathogenesis of the negative effects of opioids.

One reward-related behavior that develops during adolescence is social behavior [22–25]. In rodents, social behavior milestones during adolescence include a reduction in juvenile social play behavior and an increase in copulatory behavior, both of which need to occur for normal adult social behaviors to be achieved [26, 27]. Critical periods mark developmental periods of increased vulnerability to adverse experiences such as substance use [28–30]. We previously demonstrated that developmental changes in the nucleus accumbens (NAc) reward region regulate social play in rats during sex-specific adolescent periods: early to mid-adolescence in males (postnatal day (P)30-40) and pre-early adolescence in females (P20-30) [26]. We thus hypothesized that the developmental stage of morphine exposure will differentially impact social behavior development in each set such that: (1) morphine administered during the female critical period will result in adult sociability deficits in females, but not males, while (2) morphine administered during the male critical period will result in adult sociability deficits in males, but not females. We restricted morphine administration to either the female or the male adolescent critical periods in rats, and then measured their sociability in adulthood. We found that morphine exposure during the female critical period primarily resulted in deficits in sociability in females, while morphine exposure during the male critical period resulted in deficits in sociability primarily in males. However, depending on the test performed and the social parameter measured, social alterations could be found in both sexes during morphine exposure at either adolescent stage. These data indicate that when drug exposure occurs during adolescence, and how the endpoint data are measured, will play a large role in determining the effects of drug exposures on social development.

## 2. METHODS

### 2.1. Animal model

Adult male and female Sprague-Dawley rats were purchased for breeding (Envigo) and group housed with ad libitum access to food and water in cages with cellulose bedding changed twice weekly. Colonies were maintained in a 12:12 light:dark cycle (lights on at 07:00) in a temperature and humidity-controlled room. Litters were culled to a maximum of 12 pups per dam between P2 and P5. At P21 pups were weaned into same-sex pairs. Rats were housed in the Animal Resources Facility in Albany Medical College and all animal work was approved by the Institutional Animal Care and Use Committee at Albany Medical College.

### 2.2. Morphine exposure

Rats were assigned to one period of morphine exposure, either P23-27 (female critical period) or P31-35 (male critical period) and further grouped to receive either vehicle or morphine sulfate (Mallinckrodt, H11662) subcutaneously twice daily between the hours of 08:00-11:00 (first injection) and 15:00-18:00 (second injection). Prior to morphine or vehicle exposure, rats to be exposed to morphine during the female critical period were handled for ∼5 mins twice daily on two separate days. Rats to be exposed during the male critical period were handled for ∼5 mins once daily on five separate days. During the final handling session for rats to be exposed to morphine during both critical periods, injection acclimation with vehicle was performed. Irrespective of the age exposure, morphine was injected at a dose of 3mg/kg for the first three days and 6mg/kg for the last two days (**Fig 1A**).

**Figure 1:**
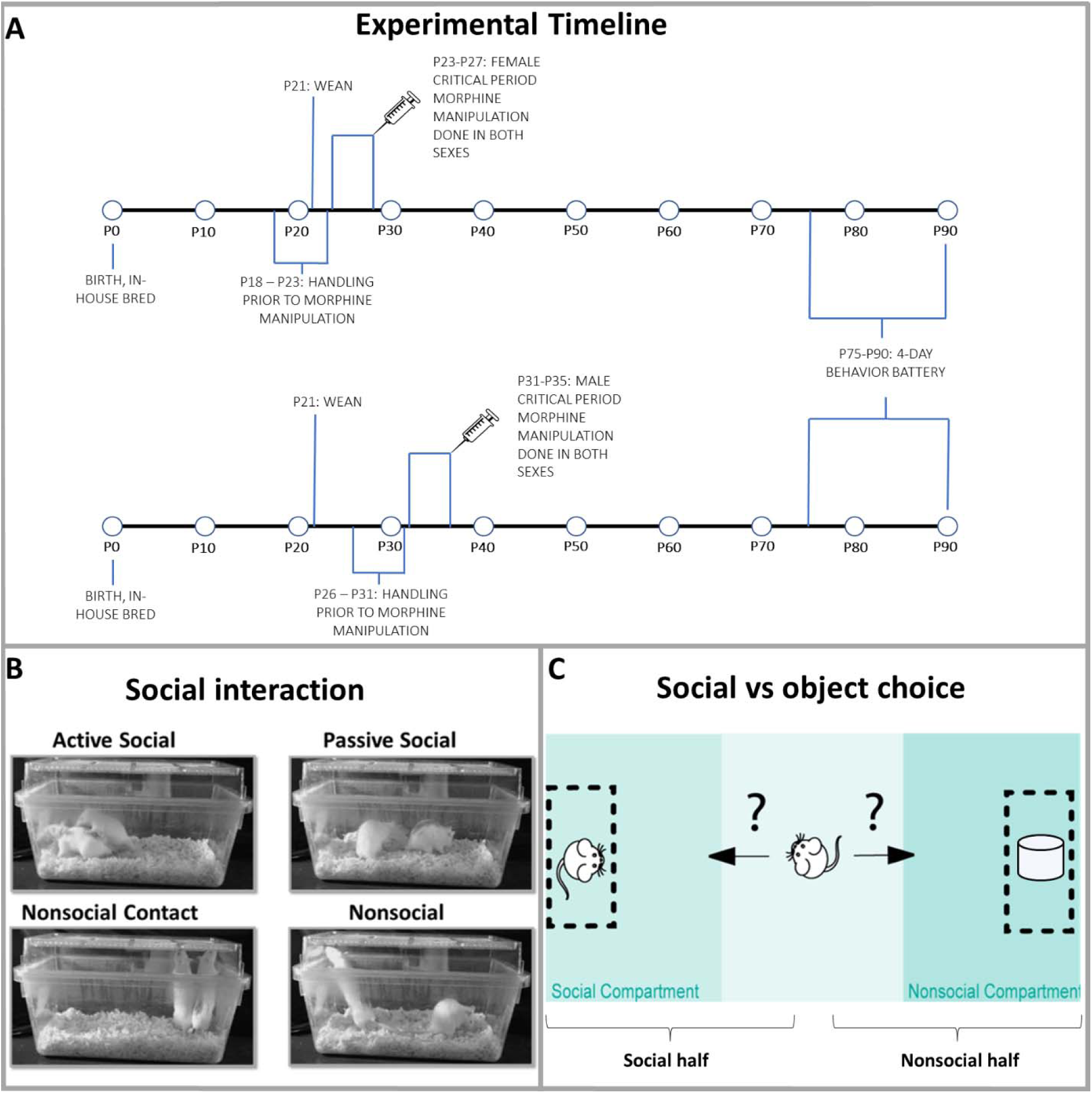
Experimental design. Visuals of **(A)** experimental timeline, **(B)** social phenotypes quantified in interaction tests and **(C)** social vs object choice assay apparatus and quantification compartments.

### 2.3. Behavioral tests

Experimental rats (P75+) and their sex- and age-matched novel conspecifics were handled for ∼5 mins on five different days in the room where all behavioral tests were to be performed. The order of behavioral tests was counterbalanced and one behavior test per day was performed. Social choice tests (which we shall refer to as social vs object choice), social interaction, and open field tests were recorded for 10 mins using ANY-Maze Behavioral tracking software. The first 5 mins of social vs object choice and social interaction tests and full 10 mins of open field were used for analysis. For the open field test experimental rats were placed in an empty open field box with overhead recording. The time spent in the center (inner zone) was automatically quantified using ANY-Maze software. For social vs object choice testing rats were first acclimated (∼10mins) in the empty three-chamber box. The three-chamber box is a behavior box with three compartments, an experimental compartment is placed in between two flanking smaller compartments where rats or objects can be placed and separated from the experimental compartment by a transparent plastic wall with several small holes not large enough for an animal to move across. Overhead video recordings of the social vs object choice tests were collected and hand coded by a blinded experimenter with Solomon Coder software (Andras Peter; solomoncoder.com) for the time spent in social exploration (zone in experimental compartment proximal to novel conspecific) and nonsocial exploration (zone in experimental compartment proximal to novel object) (**Fig 1B**). Social interaction tests were performed in empty rat housing cages. Rats were acclimated for ∼10 mins to the interaction cages prior to each experiment. Novel social interaction was performed by placing an experimental rat and a novel age- and sex-matched conspecific in the same cage for 10 mins. Familiar social interaction was performed with pair-housed cage mates re-paired for the test after a ∼20min separation. Interaction behaviors were quantified using Solomon Coder software separating out four social behavior phenotypes, 1. Active social is defined as the rat facing conspecific and physical contact with conspecific, 2. Passive social is defined as the rat facing the conspecific and no physical contact with the conspecific, 3. Nonsocial contact is defined as the rat not facing the conspecific and physical contact with the conspecific, 4. Nonsocial attention is defined as the rat not facing the conspecific and no physical contact with the conspecific. Visual depiction of the phenotypes is provided in (**Fig 1C**).

### 2.4. Statistical analysis

Data analysis was performed on Graphpad Prism version 9.5.0. Outliers were identified using the ROUT method (Q=1%). Two outliers from P23-27 familiar interaction (2 male morphine rats), 6 outliers from P31-35 familiar interaction (4 female morphine and 2 male morphine rats), 1 outlier from P31-35 novel interaction (1 male morphine rat), and 2 outliers from P31-35 social vs object choice (1 male vehicle and 1 male morphine rat) were excluded. Two-way ANOVAs were used to calculate main effects (Drug and Sex) and interaction (Drug X Sex). P23-27 and P31-35 exposure groups were performed at different times rather than simultaneously and had different handling protocols due to pre-weaning handling being required for manipulation during the female critical period. Thus, we will analyze those data separately rather than including adolescent stage as a third independent variable. If ANOVA main effect or interaction values were statistically significant, post-hoc comparisons were quantified using Sidak’s post hoc test. *p*<0.05 is defined as statistically significant.

## 3. RESULTS

### 3.1. Morphine exposure during both critical periods influences social interaction with a familiar conspecific in a sex-specific manner

To test behaviors with a familiar social partner, rats were paired with their cage mates for a social interaction test. The time spent engaged in either active social, passive social, nonsocial contact or nonsocial attention were measured. In rats exposed to morphine during the female critical period, there was an interaction of drug x sex (*F*(1,28)=4.771, *p*=0.0375) (**Fig 2D**). Post-hoc multiple comparisons tests revealed that morphine reduced time spent by females in nonsocial attention (*t*(28)=3.429, *p*=0.0038) but males were unaffected (*t*(28)=2.374, *p*=0.9932). There was a main effect of drug indicating that morphine led to rats spending less time in nonsocial attention as compared to vehicle-treated rats (*F*(1,28)=4.771, *p*=0.0375) (**Fig 2D**). In rats exposed to morphine during the male critical period, there was an interaction of drug x sex in active social interaction (*F*(1,38)=4.264, *p*=0.0458) (**Fig 2E**). Post-hoc multiple comparisons tests revealed that morphine reduced time spent by males in active social (*t*(38)=2.800, *p*=0.0159) but females were unaffected (*t*(38)=0.1778, *p*=0.9804). There was a main effect of drug that indicated that morphine increased the time spent in nonsocial attention(*F*(1,38)=7.389, *p*=0.0062) (**Fig 2H**). In addition to the drug-related effects, there were also main effects of sex. In rats exposed to morphine or vehicle during the female critical period males spend more time in active social interaction than females (*F*(1,28)=7.297, *p*=0.0116) (**Fig 2A**) and females spent more time in nonsocial attention than males (*F*(1,28)=7.028, *p*=0.0131) (**Fig 2D**). In rats exposed to morphine or vehicle during the male critical period males spent more time in active social compared to females (*F*(1,38)=4.951, *p*=0.0321) (**Fig 2E**) and nonsocial contact (*F*(1,38)=4.746, *p*=0.0356) (**Fig 2G**), while females spent more time in nonsocial compared to males (*F*(1,38)=10.96, *p*=0.0020) (**Fig 2H**).

**Figure 2:**
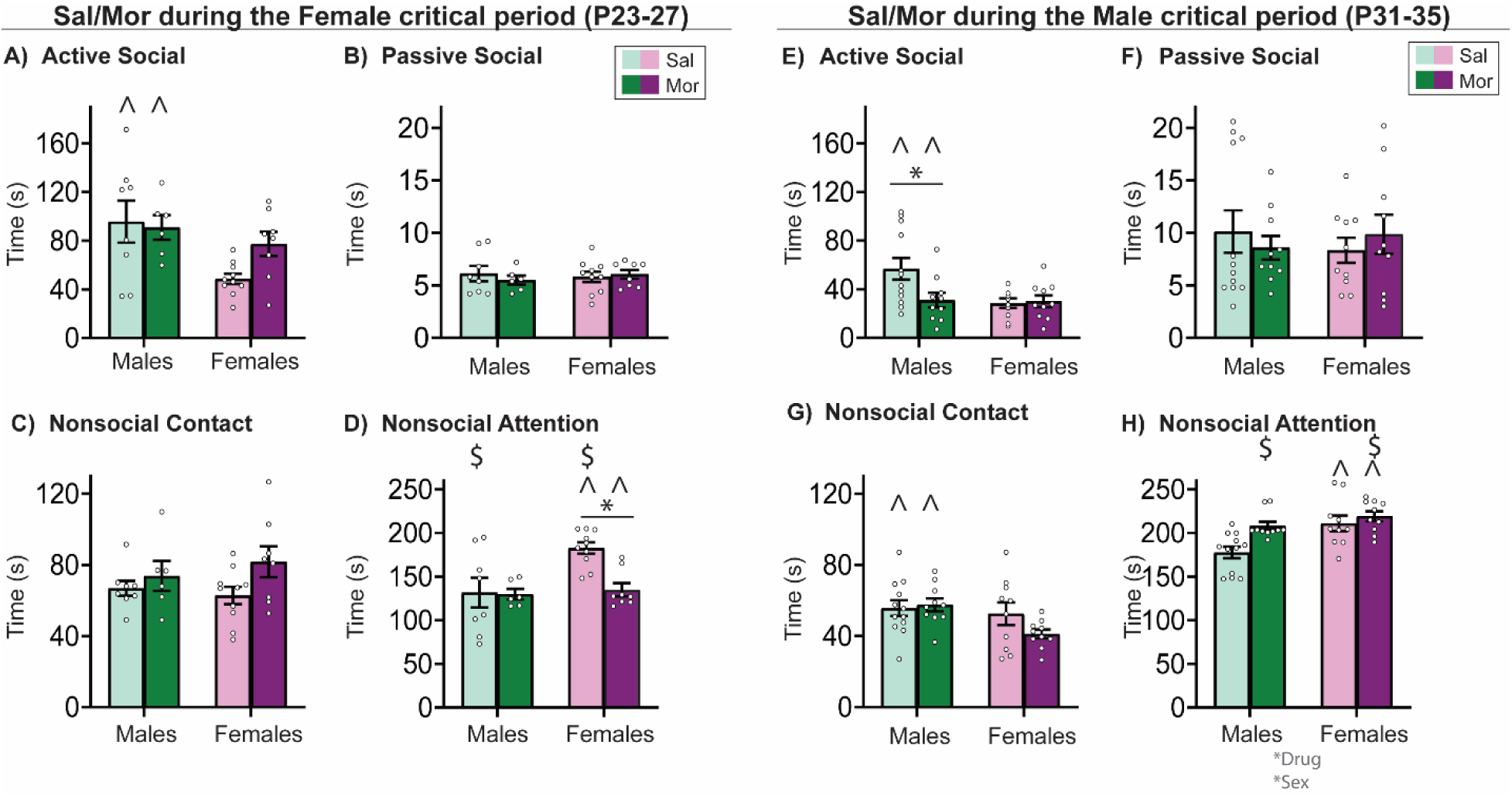
Morphine exposure during the female critical period for social development decreased nonsocial attention in females only, while morphine exposure during the male critical decreased active social interaction and increased nonsocial interaction in males only with a familiar social stimulus. (**A**) Male rats spent more time in active social compared to females. There were no statistically significant results in (**B**) passive social and, (**C**) nonsocial contact. (**D**) Female rats spent more time than males in nonsocial attention and morphine decreased the time rats spent in nonsocial attention. (**E**) There was an interaction of drug X sex in active social which showed that morphine reduces active social in males only. (**F,G**) There were no statistically significant results. (**H**) Female rats spent more time than males in nonsocial attention and morphine increased the time rats spent in nonsocial attention. Histograms depict average ± SEM. *p*<0.05 is defined as statistically significant. * is significant interaction of drug x sex post hoc test, $ is significant main effect of drug, ^ is significant main effect of sex. *n* = 8-12 rats/group.

### 3.2. Morphine exposure during the male, but not female, critical period alters social vs object choice in males only

The data thus far generally supported our hypothesis that morphine exposure during the female critical period for social development would primarily impact female social development, while morphine exposure during the male critical period would primarily impact male social development. However, familiar interaction tests are just one type of social behavior. We next sought to better understand how morphine exposure would impact novelty-directed social behaviors. In social vs. object choice tests, there was no evidence that morphine exposure during the female critical period influenced the social vs object choice in males nor females (**Fig 3A-D**). In rats that were exposed to morphine during the male critical period, there was an interaction between drug and sex in the total amount of time spent in the social (*F*(1,28)=6.782, *p*=0.0146) and nonsocial halves (*F*(1, 28)= 6.782, *p*=0.0147) of the arena. Post-hoc multiple comparisons tests revealed that morphine exposure during this adolescent stage reduced the time that males spent in the social compartment (*t*(28)=2.374, *p*=0.0488) without impacting females (*t*(28)=1.305, *p*=0.3642). Data also showed that males spent more time in the nonsocial half than females (*F(1,28)=11.65, p=0.0020*) (**Fig 3E**). To determine whether the total exploration of either stimuli was also altered, we next performed statistics on the time directly exploring the social and nonsocial stimuli. There was a significant drug x sex interaction in time spent exploring the social stimulus (*F*(1,28)=6.755, *p*=0.0147), and post-hoc tests revealed that morphine decreased social exploration in males only (*t*(28)=2.374, *p*=0.0354; (**Fig 3G**)). However, there was no change in object exploration despite P31-35 morphine-treated males spending more time in that half of the arena (*F*(1, 28) = 0.9488, *p*=0.3384; **Fig 3H**). In addition to the drug-related effects, there were also main effects of sex. Females spent less time in the social half (*F*(1,28) = 11.65, *p*=0.0020), and more time in the nonsocial half, as compared to males (*F*(1,28)=11.65, *p*=0.0020) (**Fig 3F**). Additionally, females spent more time in nonsocial exploration relative to males (*F*(1, 28) = 19.67, *p*=0.0001). These differences were not observed when vehicle or morphine injections were delivered during the female critical period (P23-27), which we will discuss further in the Discussion.

**Figure 3:**
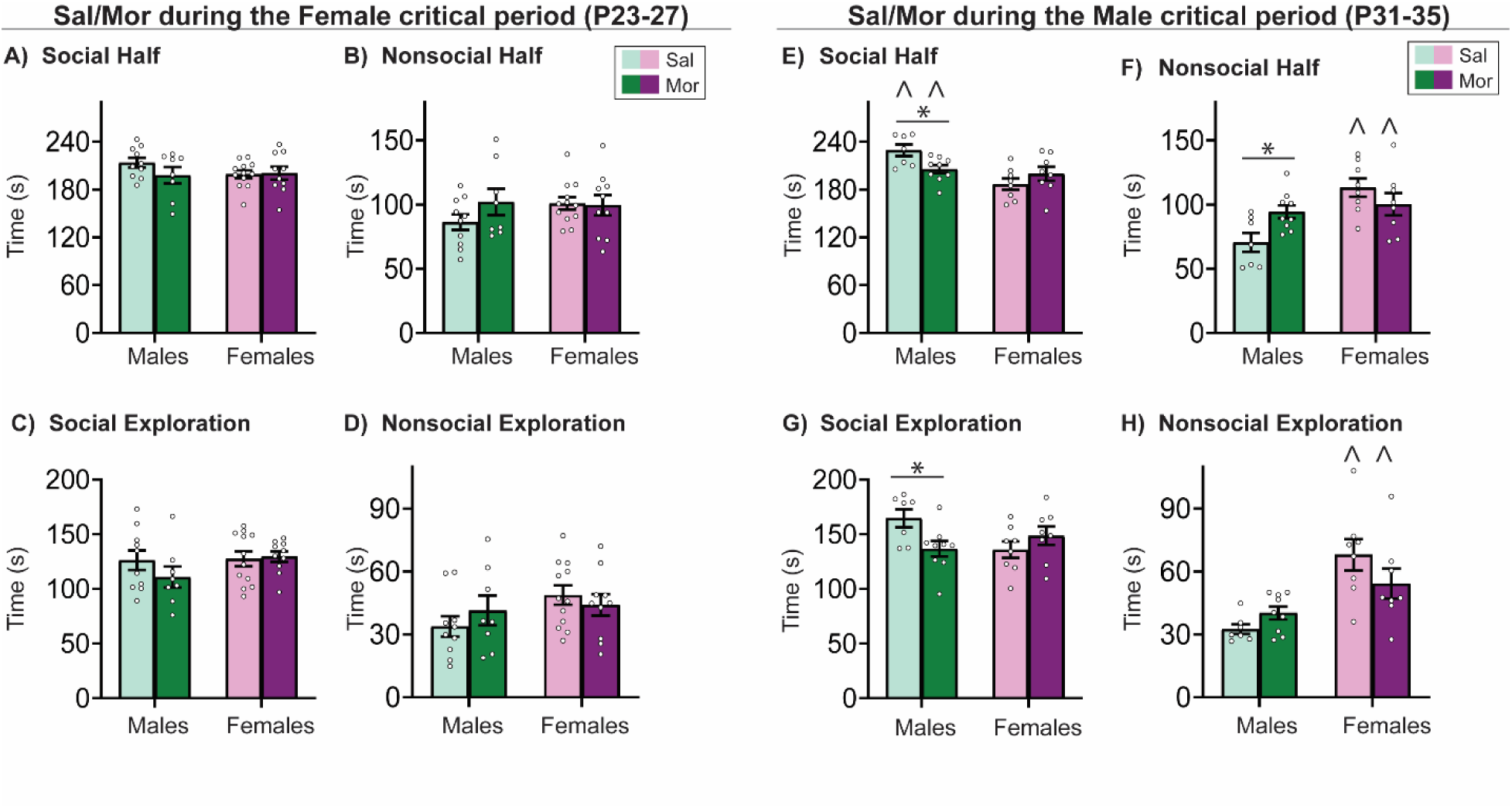
Morphine exposure during the male, but not female, adolescent critical period for social development reduces social exploration in a Social vs. Object Choice test in males only. (**A-D**) There were no statistically significant results in P23-27 data (**E**) There was an interaction of drug X sex in P31-35 in the social half. Morphine decreased time spent in the social half in males only. Males spent more time in the social half than females. (**F**) There was an interaction of drug X sex in P31-35 in the nonsocial half. Morphine increased time spent in the nonsocial half in males only. Females spent more time in the nonsocial half than males. (**G**) There was an interaction of drug X sex in P31-35 in social exploration. Morphine decreases the time spent in social exploration in males only. (**H**) Females spent more time exploring the nonsocial stimulus than males. Histograms depict average ± SEM. *p*<0.05 is defined as statistically significant. * is significant interaction of drug X sex post hoc test, ^ is significant main effect of sex. *n* = 8-12 rats/group. *n* = 8-12 rats/group.

### 3.3. Morphine exposure during the male, but not female, critical period alters social interaction with a novel conspecific

To test the effect of morphine on free social interaction, social interaction tests were performed with a novel age- and sex-matched social conspecific. The time spent engaged in either active social, passive social, nonsocial contact or nonsocial attention were measured. In rats exposed to morphine during the female critical period for social development, there was no effect of morphine in either sex in any of the behavior features quantified (**Fig 4A-D**). In rats exposed to morphine during the male critical period for social development, there was a main effect of drug indicating that morphine increased the time spent in active social interaction (*F*(1,36)=4.21, *p*=0.0475) (**Fig 4E**). Other behavioral features were not influenced by morphine exposure during the male critical period (**Fig 4F-H**).

**Figure 4:**
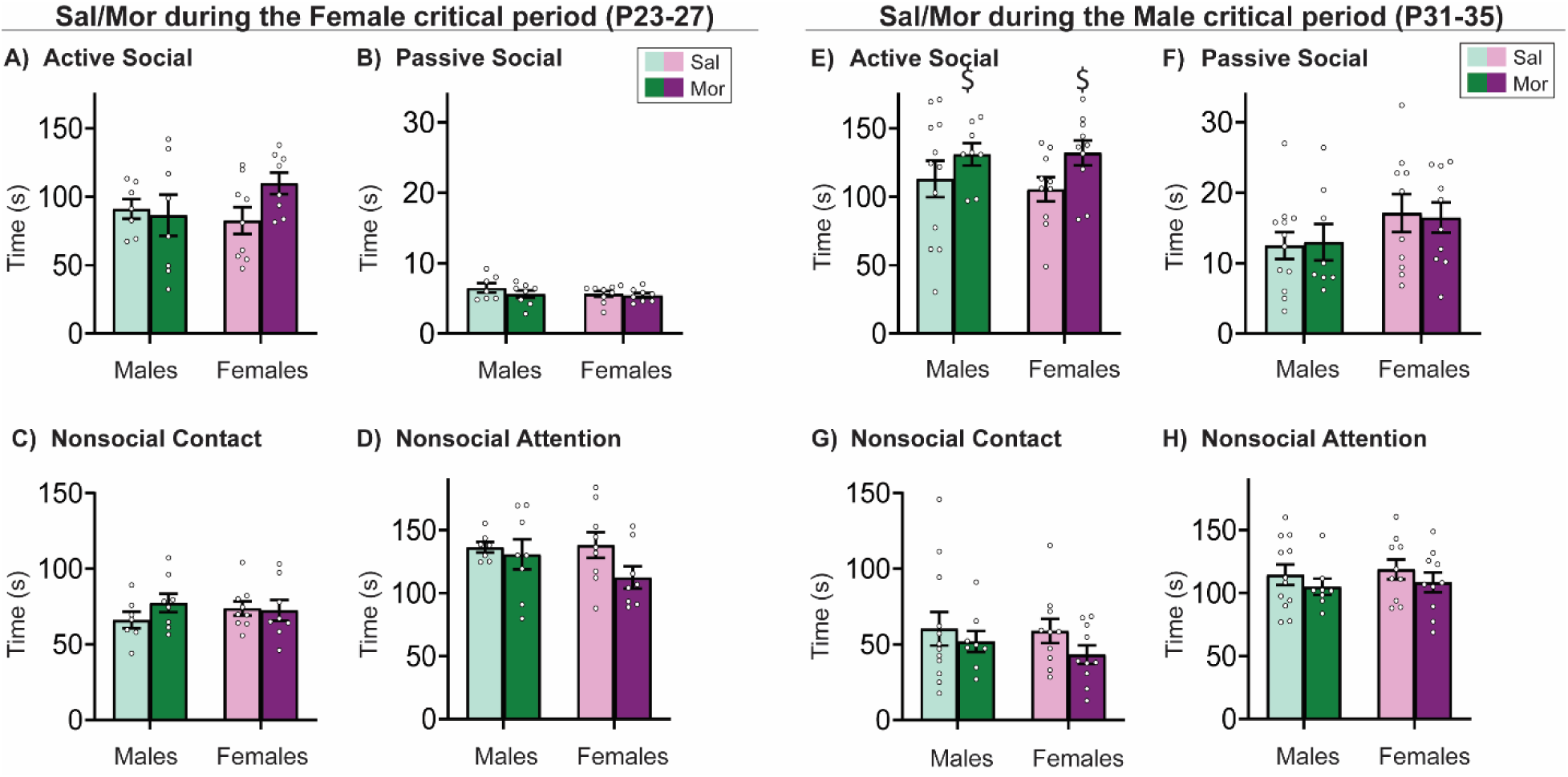
Morphine exposure during the male, but not female, adolescent critical period for social development increased active social interaction with a novel social stimulus in a sex-independent way. (**A-D**) There were no statistically significant results in P23-27 data. (**E**) There was a statistically significant effect of drug in P31-35 active social. Morphine increased the time spent in active social in both sexes. (**F-H**) There were no statistically significant results. Histograms depict average ± SEM. *p*<0.05 is defined as statistically significant. $ is significant main effect of drug. *n* = 8-12 rats/group.

### 3.4. Adolescent morphine exposure during both critical periods for social development influences free social interaction in naïve conspecifics paired with morphine exposed rats

We next tested whether social responses of the naïve conspecifics used in the novel social interaction test would vary based on whether their social partner (the experimental rat) was exposed to morphine or vehicle during the male or female critical periods. In naïve rats paired with rats exposed to morphine during the female critical period for social development there was an interaction of drug x sex in nonsocial contact (*F*(1,26)= 4.303, *p*=0.0481) (**Fig 5C**). Post-hoc multiple comparisons tests revealed that morphine exposure in the experimental rats induced naïve rats to decrease the time spent in nonsocial contact in males (*t*(26)=3.078, *p*=0.0097) but not females (*t*(26)=0.2579, *p*=0.9594) (**Fig 5C**). There was a main effect of drug indicating that morphine exposure in the experimental rats induced an increase in the conspecific’s time spent in active social (*F*(1,26)= 8.957, *p*=0.0060) (**Fig 5A**) and a decrease in time spent in nonsocial contact (*F*(1,26)= 5.887, *p*=0.0225) (**Fig 5C**). In naïve rats paired with rats exposed to morphine during the male critical period for social development there was an interaction of drug x sex in nonsocial attention (*F*(1,34) = 5.528, *p*=0.0247) (**Fig 5H**). Post-hoc multiple comparisons tests were not significant.

**Figure 5:**
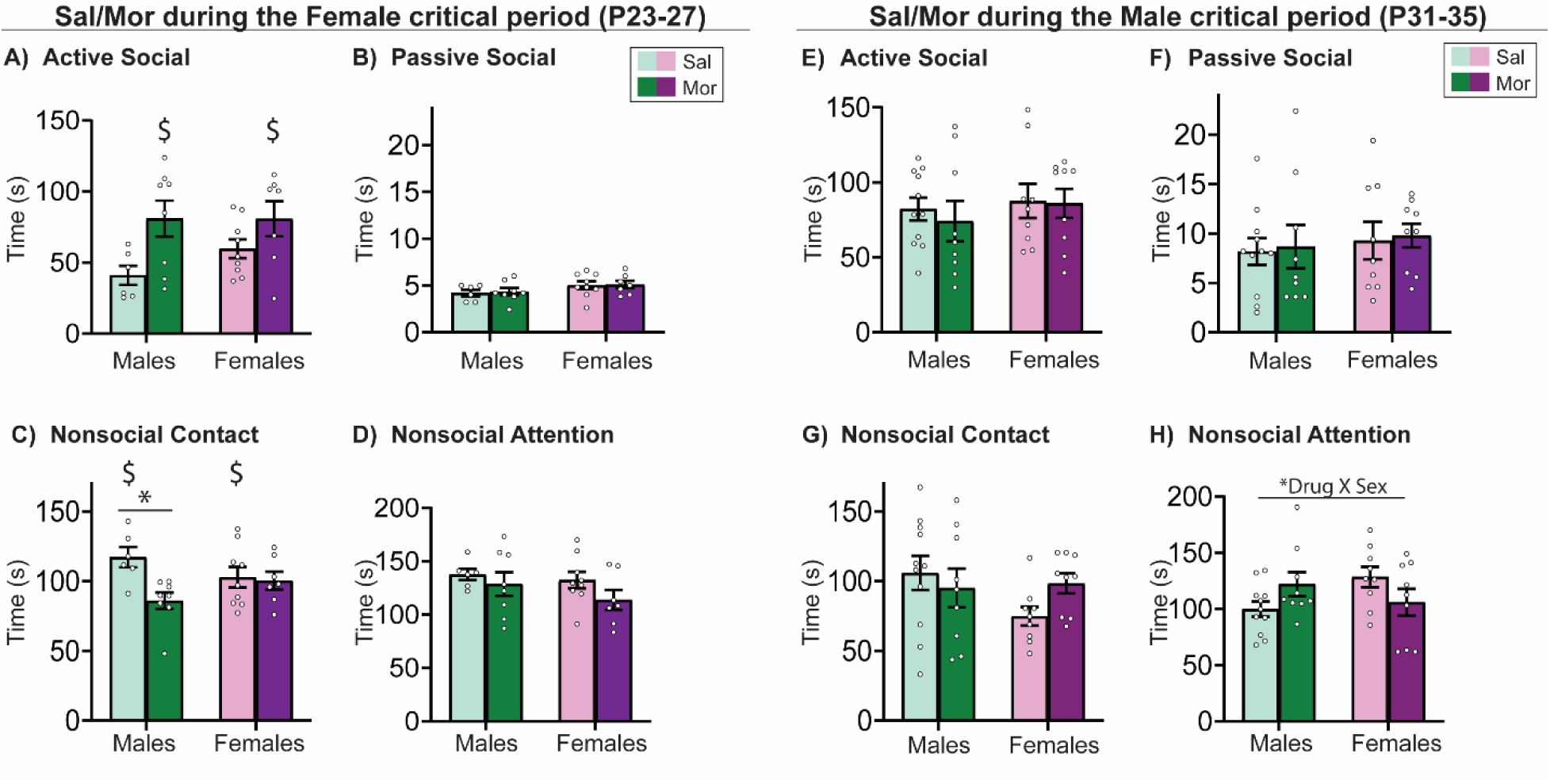
Interaction with morphine exposed rats reveals increased active social in both sexes when paired with rats exposed to morphine during the female but not male critical period of social development. (**A**) Naïve rats spent more time with morphine exposed rats of both sexes are compared to vehicle treated rats. (**C**) Male naïve rats spend less time with morphine exposed rats in nonsocial contact and morphine reduced overall time spent in nonsocial contact as compared to vehicle exposed rats (**B,D**) There were no statistically significant results. (**E-G**) There were no statistically significant results (**H**) There was an interaction of drug X sex in the amount of time spent by naïve rats in nonsocial attention indicating both sexes of naïve rats responded to morphine exposed rats in a different manner. Histograms depict average ± SEM. *p*<0.05 is defined as statistically significant. * is significant interaction of drug X sex post hoc test, $ is significant main effect of drug. *n* = 8-12 rats/group. *n* = 8-12 rats/group.

### 3.5. Morphine exposure during adolescent critical periods for social development does not influence anxiety-like behavior in open field

There was no main effect of drug after morphine exposure during the male and female critical periods indicating that morphine does not influence the time spend in the inner zone of the open field. Open field tests whether morphine influenced anxiety-like or exploratory behavior after adolescent morphine exposure. This sheds light on whether the effects in the social tests result from altered anxiety-like or exploratory phenotype, and not because of differences in social development. After morphine or vehicle exposure during the female critical period, we found a main effect of sex in both the time rats spent in the inner zone (*F*(1,36)=12.79, *p*=0.0010) of the open field with males spending more time exploring the inner zone than females (**Supplemental Fig 1A**). Likewise, when morphine or vehicle manipulation was performed during the male critical period there was a main effect of sex in the time the rats spent in the inner zone (*F*(1,42)=19.50, *p*=0.0001) with males spending more time exploring the inner zone than females (**Supplemental Fig 1B**).

## 4. DISCUSSION

Our data demonstrate that even a short-term morphine exposure during adolescence can persistently influence the phenotypes observed in social behavior. Moreover, how one assesses social behavior can also impact whether a social phenotype will be observed after adolescent morphine exposure.

Herein we discuss social behavior under the construct of sociability. Sociability is the tendency of individuals to associate with other individuals. This behavioral construct excludes behaviors that are primarily influenced by reproductive- and parental-related behaviors [31–33]. Sociability can be further broken down into the identification, classification, integration, and response to social cues [34] and is among the most complex sets of behavioral features controlled by the central nervous system [35]. Our study focuses on changes in sociability in the context of rat models of social vs object choice and free social interaction tests. In social vs object choice, we define an increase in sociability as an increase in the time spent in social exploration and/or the social half. Interaction tests probed two aspects of sociability: (1) social touch is rats having physical contact with conspecifics [36–39] and is quantified in active social interaction and nonsocial contact, and (2) Visual social attention is rats having conspecifics in direct gaze [40–43] and is quantified in active and passive social interaction. Active social interaction is a mixed social phenotype containing both aspects of sociability while nonsocial attention is the absence of any social aspects quantified. Thus, we define an increase in sociability as an increase in time spent in active social, passive social or/and nonsocial contact. Finally, we found that morphine exposure did not alter anxiety-like or exploratory behaviors in the Open Field test (**Supplemental Fig.1**). Thus, we interpreted results based on the assumption that deficits in social tests resulted from changes in sociability and were not affected by anxiety-like phenotype.

### 4.1. Sex-specific vulnerability of social development to morphine during adolescence is dependent on adolescent period

Epidemiological data in humans indicates that acute exposure to opioids during adolescence increases the risk of psychiatric illness and substance use disorder later in life [13–18]. Developmental differences in response to opioids are documented in behavior [44], pain and hyperalgesia [45], glia [46] and neurons [47]. We demonstrate that developmental stage and sex also play a role in the vulnerability to opioid exposure. This is an important link built on the knowledge that males and females adolescent development occurs at different periods, with females generally moving into puberty earlier than males [26, 48–53] including social development [26, 54, 55]. Therefore, we expected that morphine-induced sociability deficits would differ by period of exposure, and that female deficits would be restricted to morphine exposure during the female critical period and vice versa for males. The reality was more complex, which we will discuss further below. Discriminating sex and period specific opioid use vulnerabilities during adolescent social development will inform best practices around opioid prescriptions for adolescents which impair social behavior later in life. However, unexpectedly, these two injection periods also appeared to be differentially modulated by vehicle injections. Injection stress has been reported to alter normal development in rodent models [56, 57]. We did not have an injection-naïve group for comparison to directly test injection stress, however, we report that period of injection stress results in different levels of sociability that prevented us from analyzing the data using a statistical approach to include period of morphine exposure (3-way ANOVA: drug x sex x age). Thus, these adolescent periods may be vulnerable not just to morphine or injection stress, but to many different stressors. The Adolescent Brain Cognitive Development (ABCD) study is a large long-term study of juvenile neurodevelopment [58–60]. The ABCD study may reveal grey and white matter changes that are associated with adolescent adverse experiences such as the effects of acute opioid exposure. A debate about the cumulative versus dimensional buildup of long-term effects from adverse adolescent experiences is ongoing [61–63]. While there is agreement that multiple adverse experiences across development lead to worsening effects, whether these effects build up by worsening of specific behavior features or whether different adverse experiences alter different behavior features are in the early stages of study [64]. It is plausible that adolescent opioid exposure primes individuals for worsening social development and subsequent adverse experiences cause progression of deficits. It will be important to determine in future studies whether different addictive substances or other stressors impact social development in similar or different ways between the sexes. An integrated model of unexpected stress after adolescent morphine exposure during the male and female critical period may answer the question of the sex and period specific vulnerabilities followed by additional adverse experiences.

### 4.2. Social abnormalities depend on how you assess social behavior

Social context influences the type of social response [22, 65, 66]. For example, Veenema *et al* showed that inhibiting vasopressin receptors had no effect on social interaction in a novel cage but social interaction in home cage was *increased* by inhibition of vasopressin receptors [67, 68]. This result indicates that the same social circuit can respond differently to different contexts of the same behavior.

Familiar social interaction assesses the quality of free social behavior with a familiar conspecific. Males, but not females, exposed to morphine during the male critical period of social development had decreased sociability (reduced active social interaction) indicating a male-specific social developmental vulnerability induced by morphine at this stage of adolescence (**Fig 2E**). Conversely, morphine exposure during the female critical period increased sociability (reduces nonsocial behaviors) compared to vehicle groups in females, but not males. Although familiar interaction tests are infrequently used in the literature, we propose they make a strong addition to a social behavioral battery. Familiar social relationships like families fill a core and often life-long social role in one’s life [69, 70]. In our studies, rats were pair housed with a sex-, age- and treatment group-matched rat, and thus these rats had a matched history of morphine or vehicle exposure. Hence, if morphine exposure did induce social deficits, we expected these to be magnified due to the presence of at two socially deficient rats. Substance use disorder is affected by worsening of both familial and broad social bonds [71], our results suggest that even an acute exposure leads to lasting social deficits. Thus, with respect to models of drug exposures, we argue that familiar interaction tests are an essential complement to tests with novel social stimuli and may be more sensitive to subtle differences in social behaviors.

The social vs object choice test assesses the natural proclivity for rats to prefer novel social stimuli over novel nonsocial stimuli [72] and is a frequently used and simple test of sociability in rodents. Unlike the familiar interaction test, neither male nor female rats exposed to morphine during the female critical period for social development had any changes in sociability in this test (**Fig 3A,C**). Similar to the familiar interaction test, males, but not females, exposed to morphine during the male critical period for social development had reduced sociability (**Fig 3E,G**).

Novel social interaction test assesses the quality of free social behavior with a novel conspecific [73, 74]. To our surprise, there was very little change in novel social interaction in the experimental rats (those that received morphine or vehicle during adolescence). Morphine exposure during the female critical period for social development had no sociability deficits (**Fig 4A-D**), while there was a main effect of morphine exposure during the male critical period, resulting in increased sociability (increased active social interaction) that appeared to be present in both sexes (**Fig 4E**).

There are few instances in the literature in which the conspecific’s social behavior is coded alongside the experimental target’s behavior. Sociability is a bidirectional process where two or more individuals engage in an association [75–77]. Research in humans indicates that people tend to spend more time interacting with neurotypical persons [78–80], this can be among the reasons people with psychiatric illness or substance use disorder are more likely to have insufficient social connections that get worse with time and disease progression [81–83]. Understanding the responses naïve individuals have to individuals that have social deficits is therefore an important aspect to model. Thus, we decided to code both sets of social behavior in case there was an altered social response in conspecifics due to the adolescent morphine/vehicle history of their social partners. Our data reveal that conspecific, non-manipulated rats do indeed respond differently depending on whether rats were exposed to morphine during adolescence (**Fig 5**). In fact, these differences were far more robust in the conspecifics than in the experimental animals to which they were responding. Because there are few overt social changes in the experimental rats in this test (**Fig 4**), these data suggest that social partner rats can identify differences in behavior caused by adolescent morphine exposure, which in turn elicits different behavioral responses. Interestingly, analyzing the behavior of the naïve conspecific rats may shed light on deficits that are beyond the limitations of our study design. What the conspecifics are responding to is unclear at this point, however, we speculate the possibility of olfactory, vocal, and rodent body language signals that we were not measuring in the current studies [84–86]. These more covert social cues will be important to assess in future studies. Regardless, it is important to distinguish effects of opioids in different spatial and social contexts because therapeutic approaches could be optimized by adequate social environment cues, an idea supported by work indicating that volitional abstinence from addictive substances can be induced by social interaction [87–89].

### 4.3. Importance of social function in substance use disorders

There is extensive literature acknowledging that addictive substance use during adolescence disrupts normal adolescent development [90, 91]. Adolescents are more likely than adults to initiate first time drug use [92], and the older the age of first use the less likely an individual develops substance use disorder [93, 94]. These effects may be in part be driven by the increase in peer-centered decision making [95] that occurs during adolescence. Male and female social development occurs on different timelines [96, 97].

Decreased sociability increases the likelihood of developing substance use disorder later in life [98, 99] raising an interesting question of whether our morphine model increases the likelihood of our rats developing addiction-related behaviors if exposed to voluntary drug use at a later point. Volitional sociability reduces the risk of developing addiction-related behaviors in rodents [88]. This stresses the importance of sociability in prevention and recovery from addiction-related syndromes in the clinic [100, 101].

Social influence is an important component of sociability because individuals respond to the social behavior of others especially when interacting with individuals from different levels of social hierarchies [102–104]. In rodents the behavior of a dominant male will influence the behavior of a submissive male [105, 106]. Opioids disrupting normal social development could modify social influence causing the exacerbation of abnormal social behaviors and social transfer to others as we report in sociability with a familiar conspecific (**Fig 2**) and influence the responses of naïve strangers to abnormal individual such as in (**Fig 5**). This raises the question of whether there is a component to the human epidemiological data that found increases in the incidence of psychiatric illness in adolescents after acute opioids that can be explained by changes in responses from social partners as a response to opioid induced social deficits in patients. This would be a challenging effect to isolate in clinic, however, one necessary to consider.

### 4.4. Potential mechanisms

The pharmacokinetics and pharmacodynamics of the neurobiology of morphine are known and result in the binding to opioid receptors on inhibitory neurons that synapse onto dopaminergic neurons in the ventral tegmental area. Inhibiting these inhibitory neurons leads to a disinhibition of the dopaminergic neurons which project into the nucleus accumbens causing an increase in dopamine transmission [47, 107]. Generally, in opioid models of rodents, opioid receptor agonists are observed to cause social aversion [108] some studies suggesting this effect is regulated by noradrenaline and serotonin signaling [109, 110]. However, Piccin and Contarino observed that withdrawal from chronic morphine exposure in adult mice increased social approach [111].

The adolescent developmental periods examined herein correspond to sex-specific periods of synaptic remodeling of the nucleus accumbens reward (NAc) region via synaptic pruning [26]. Synaptic pruning in this region is mediated via the phagocytosis of synapses or synaptic proteins by microglia [26, 112–114]. Adolescent social development in males and females regulated by the nucleus accumbens (NAc) requires synaptic pruning and occurs earlier in females P20-28 than males P30-38. Male synaptic pruning is aimed at downregulating Dopamine receptor 1 (DR1) positive synapses by tagging with complement factor 3 (C3), an “eat me” signal for microglia phagocytosis. In females, synapses are tagged by C3 for pruning but the target(s) for pruning on synapses are unknown [26]. When synaptic remodeling is blocked during adolescence social deficits result [26, 115, 116]. This led us to speculate that morphine exposure would influence the disruption of these developing social circuits causing lasting behavioral deficits. The effect of morphine on developing social circuits is currently unexplored. However, the answer to the question of the effect of morphine on microglia the cells type that performs synaptic pruning there are some hints. We have shown that morphine modulates phagocytosis of microglia *in vitro* through activity on toll-like receptor 4 (TLR4) [117], and thus we predict that morphine increases pro-phagocytic neuroinflammatory profile of microglia via TLR4 during sex specific critical periods causing increased pruning and sociability deficits in adulthood.

### 4.5. Limitations and future directions

The golden standard in rodent drug studies is self-administration [118, 119]. This assumes that addictive substances reinforce subsequent use [120]. Self-administration models offer the choice of the addictive substance or a non-drug reinforcer. This technique has been used to demonstrate that volitional social behavior inhibits drug addiction [88], supporting our prediction that social development is important in substance use disorder. Our model of involuntary morphine exposure while limited to answering questions on addiction-like behavior is a robust model of the clinical juvenile pain management setting of acute morphine exposure [121, 122].

Additionally, due to the early age of the critical periods (P23-27 and P31-35) it is impossible to train juvenile rats for self-administration. However, involuntary morphine exposure allows us to probe earlier timepoints for different critical periods that may be lastingly impacted by morphine exposure.

Morphine-induced exacerbated synaptic pruning is what we predict to be the main driver of the deficits we report, but we cannot claim exclusivity of this mechanism to explain these deficits. We control morphine exposure temporally increasing our probability of disrupting normal synaptic pruning processes in the manner we predict. A causal study spatially modulating synaptic pruning to normal levels in the NAc prior, during and after the critical periods, and resulting in a rescue of normal sociability during adulthood would answer this question. Validation via causal mechanism will make translational implications of our work more robust because it will give both behavioral and mechanistic predictions.

The social decision-making network is comprised of both the social behavior network and the mesolimbic reward system [123], it regulates sociability [124] and is implicated in drug-related behaviors [125]. Drug-related behaviors are primarily influenced by the integration of multiple brain networks into an addiction network [126] and brain regions of the social decision-making network are among them [127]. We appreciate the complexity involved in sociability and the implications for drug-related behaviors and propose that our study opens the door to understanding the contributions of vulnerabilities of juvenile drug exposures during critical periods for social development.

### 4.6. Conclusion

In summary, we characterized long-lasting opioid-induced social deficits that depend on period of exposure and sex. The context of social behavior revealed different types of deficits and the period of opioid exposure and the resulting effects by sex hint at the underlying mechanisms causing the deficits. The direction of deficits depending on sex is not as clear as our original hypothesis, thus our reformed hypothesis is that morphine induced deficits are period specific depending on behavioral task and can be sex specific or sex neutral. These results suggest a complex interplay of different sociability features disrupted in different ways by opioids leading to different social deficits.

## Acknowledgements

This work was supported by the National Institutes of Health R01DA052889 and R03AG07011 to AMK and Albany Medical College Start-up funds to AMK.

## Author Declarations

DNK and AMK designed the experiments. DNK, CF, and JMK performed the experiments. DNK, ELE, and AMK analyzed the experiments. DNK, ELE, and AMK wrote the manuscript. All authors edited the manuscript. The authors declare no conflicts of interest.

## SUPPLEMENTAL FIGURES AND TABLES

**Supplemental Figure 1:**
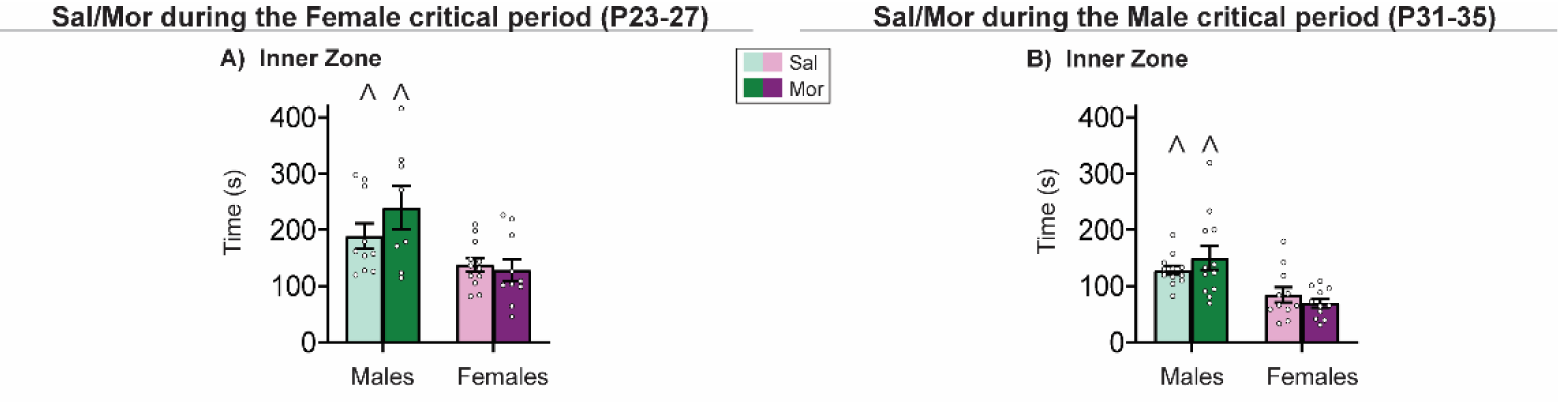
Morphine does not alter anxiety-like behavior. (**A,B**) Male rats spent more time in the inner zone that female rats regardless of period of morphine exposure. There was no regulation of anxiety-like behavior by morphine. Histograms depict average ± SEM. *p*<0.05 is defined as statistically significant. ^ is significant main effect of sex. *n* = 8-12 rats/group. *n* = 8-12 rats/group.

**Supplemental Table 1:** Detailed statistics table corresponding with Figure 2.

|  | Familiar Interaction: Statistical Main Effects |  |  |  |  |  |
| --- | --- | --- | --- | --- | --- | --- |
|  | Main effect of sex |  |  |  |  |  |
|  | (P23-P27) |  |  | (P31-35) |  |  |
|  | F (1, 28) value | P value | Outcome | F (1, 38) value | P value | Outcome |
| Active social | 7.297 | 0.0116 | M > F | 4.951 | 0.0321 | M > F |
| Passive social | 0.04847 | 0.8273 |  | 0.02196 | 0.883 |  |
| Nonsocial contact | 0.09058 | 0.7657 |  | 4.746 | 0.0356 | M > F |
| Nonsocial attention | 7.028 | 0.0131 | F > M | 10.96 | 0.002 | F > M |
|  | Main effect of drug |  |  |  |  |  |
|  | (P23-P27) |  |  | (P31-35) |  |  |
|  | F (1, 28) value | P value | Outcome | F (1, 38) value | P value | Outcome |
| Active social | 1.158 | 0.291 |  | 3.269 | 0.0785 |  |
| Passive social | 0.126 | 0.7253 |  | 1.60E-05 | 0.9968 |  |
| Nonsocial contact | 3.849 | 0.0598 |  | 1.114 | 0.2979 |  |
| Nonsocial attention | 5.483 | 0.0265 | Veh > Mor | 8.389 | 0.0062 | Mor > Veh |
|  | Interaction |  |  |  |  |  |
|  | (P23-P27) |  |  | (P31-35) |  |  |
|  | F (1, 28) value | P value | Outcome | F (1, 38) value | P value | Outcome |
| Active social | 2.232 | 0.1463 |  | 4.264 | 0.0458 | M Veh > M Mor |
| Passive social | 0.5903 | 0.4487 |  | 0.8633 | 0.3587 |  |
| Nonsocial contact | 0.8257 | 0.3713 |  | 2.204 | 0.1459 |  |
| Nonsocial attention | 4.771 | 0.0375 | F Veh > F Mor | 2.813 | 0.1017 |  |

**Supplemental Table 2:**
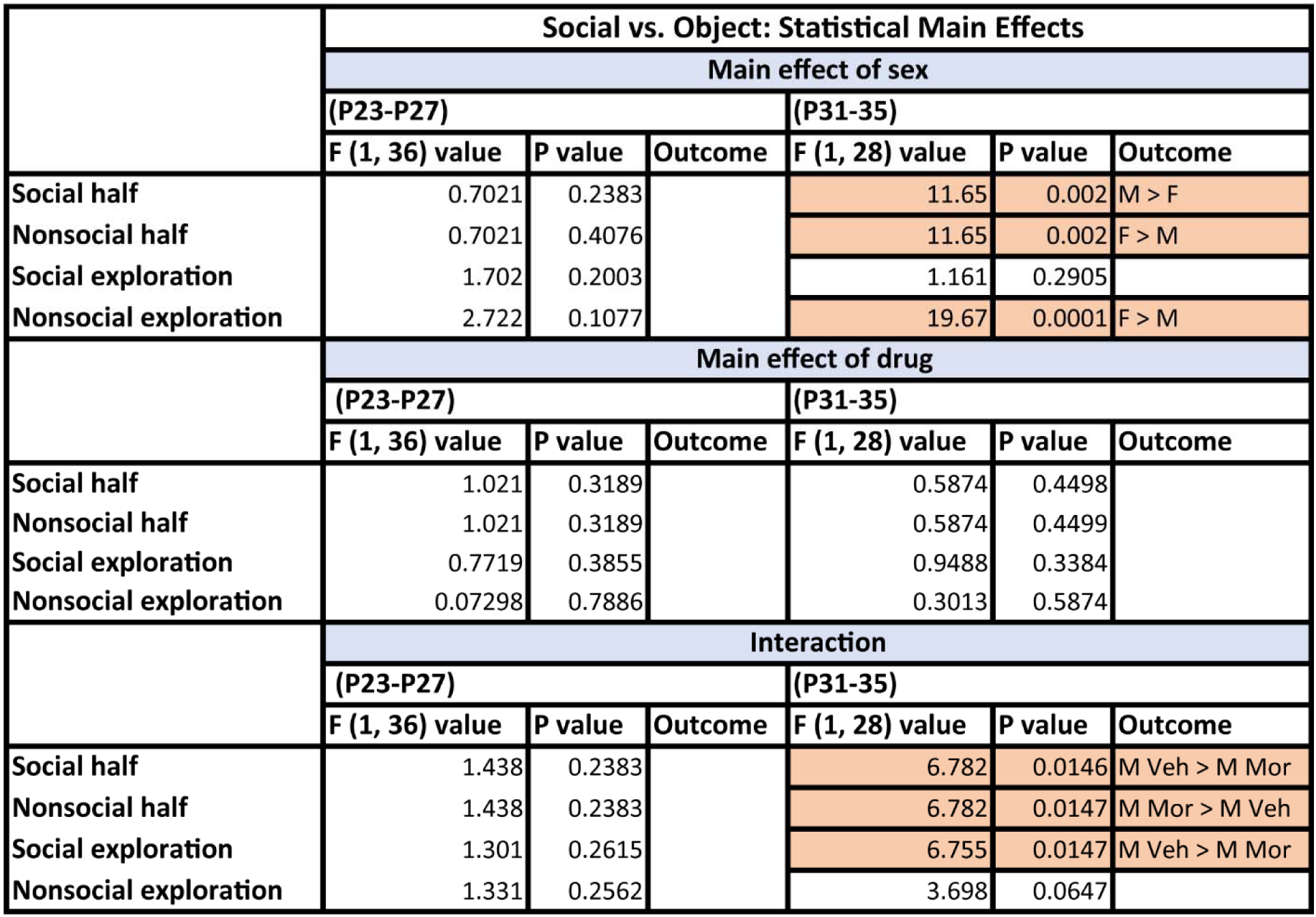
Detailed statistics table corresponding with Figure 3.

**Supplemental Table 3:**
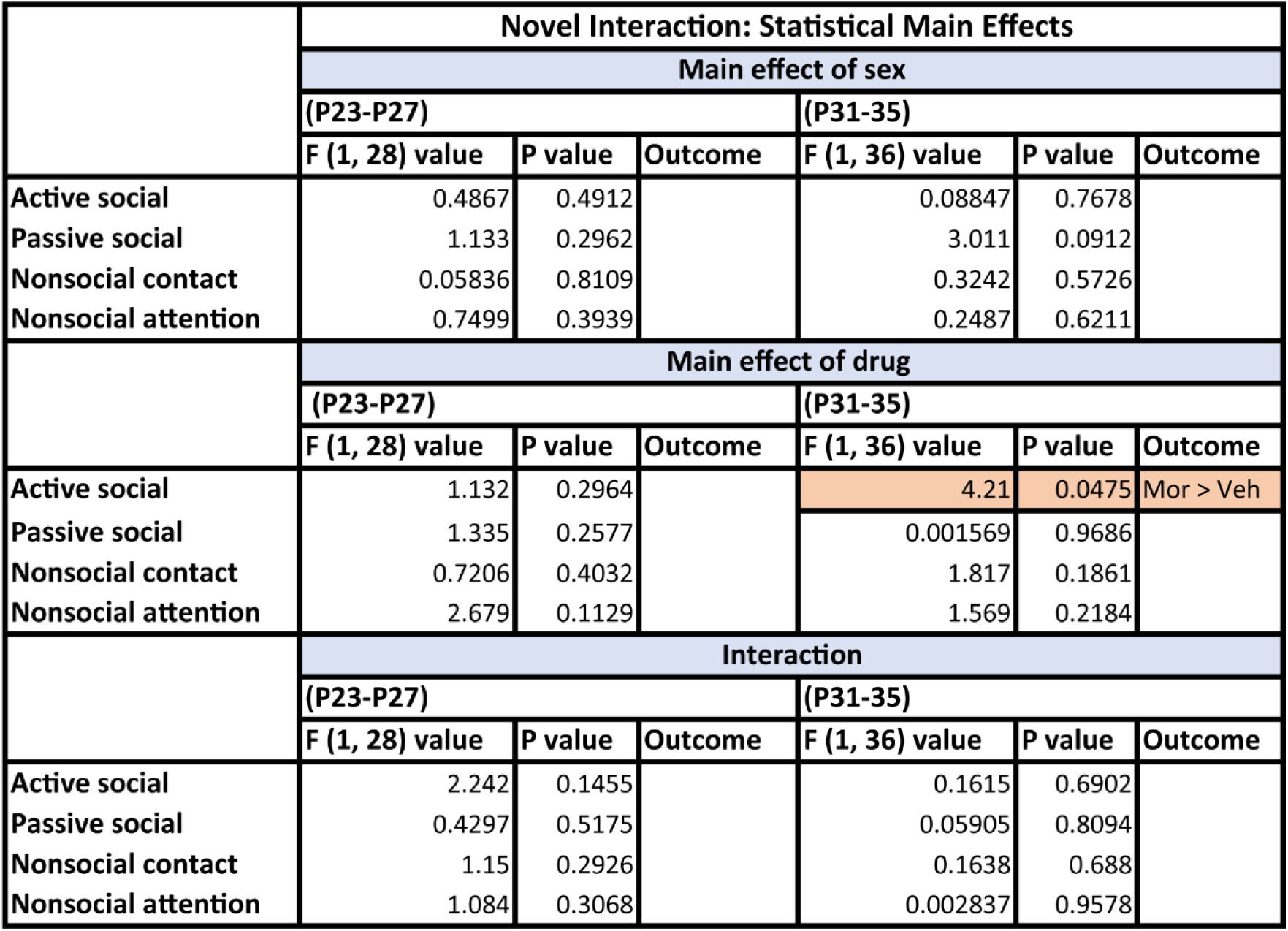
Detailed statistics table corresponding with Figure 4.

**Supplemental Table 4:**
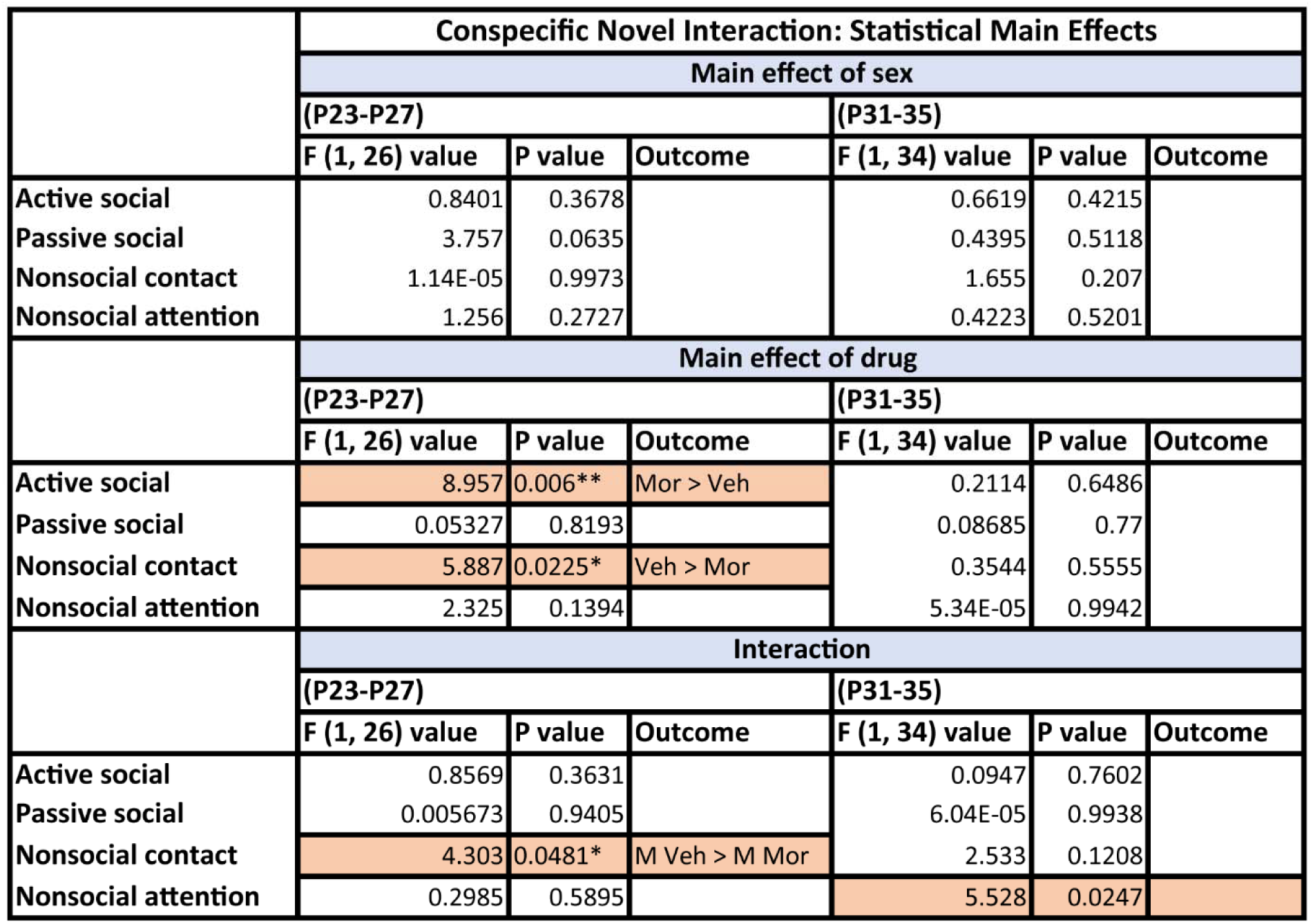
Detailed statistics table corresponding with Figure 5.

**Supplemental Table 5:**
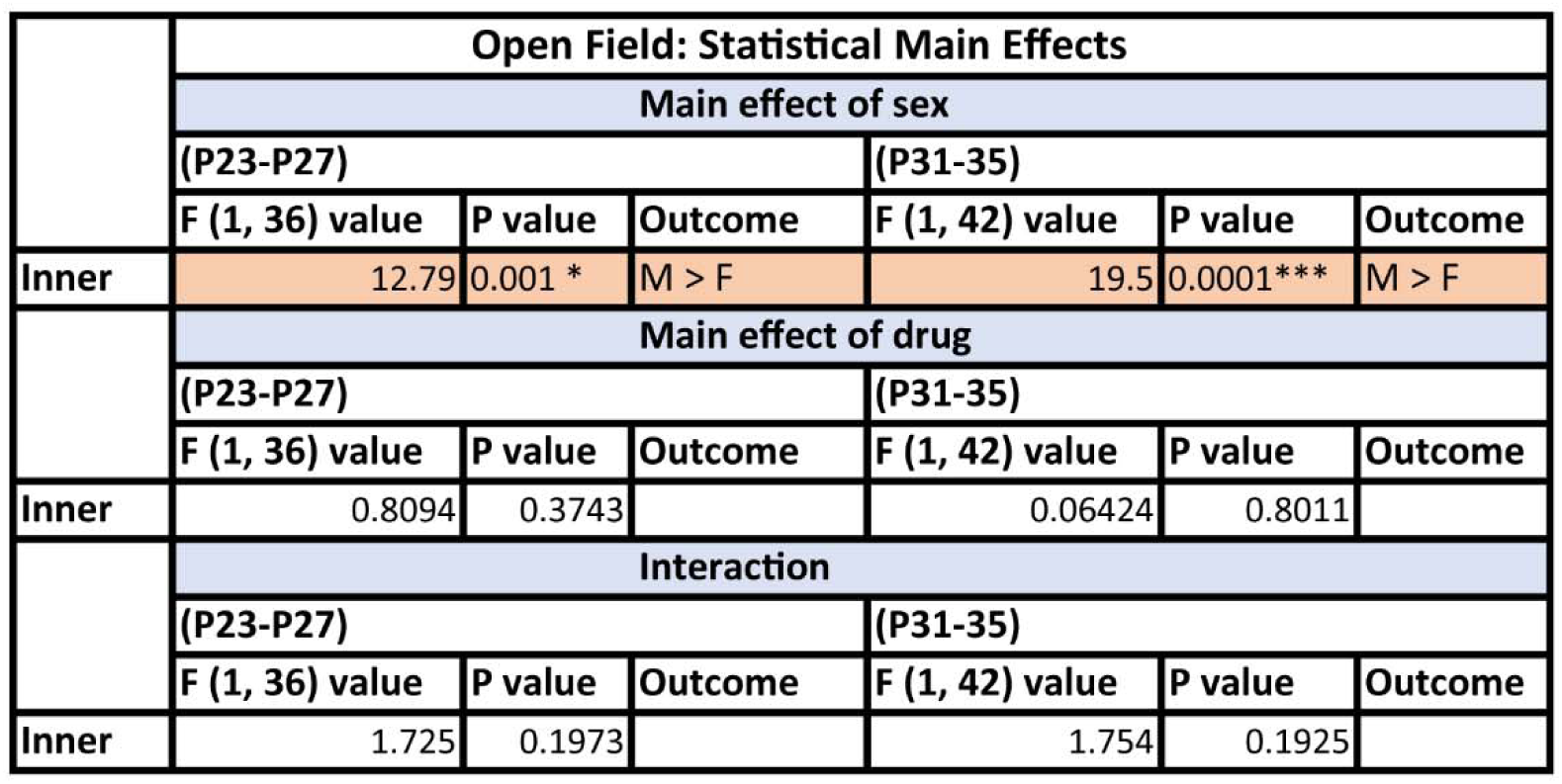
Detailed statistics table corresponding with Supplemental Figure 1.

## REFERENCES

1. Tapocik, J.D., et al., Neuroplasticity, axonal guidance and micro-RNA genes are associated with morphine self-administration behavior. Addict Biol, 2013. 18(3): p. 480–95.

2. Madayag, A.C., et al., Cell-type and region-specific nucleus accumbens AMPAR plasticity associated with morphine reward, reinstatement, and spontaneous withdrawal. Brain Struct Funct, 2019. 224(7): p. 2311–2324.

3. Lefevre, E.M., et al., Differential Patterns of Synaptic Plasticity in the Nucleus Accumbens Caused by Continuous and Interrupted Morphine Exposure. J Neurosci, 2023. 43(2): p. 308–318.

4. Salmanzadeh, H., et al., Adolescent drug exposure: A review of evidence for the development of persistent changes in brain function. Brain Res Bull, 2020. 156: p. 105–117.

5. Mooney-Leber, S.M. and T.J. Gould, The long-term cognitive consequences of adolescent exposure to recreational drugs of abuse. Learn Mem, 2018. 25(9): p. 481–491.

6. Nawi, A.M., et al., Risk and protective factors of drug abuse among adolescents: a systematic review. BMC Public Health, 2021. 21(1): p. 2088.

7. Orihuel, J., et al., ΔfJ9-Tetrahydrocannabinol During Adolescence Reprograms the Nucleus Accumbens Transcriptome, Affecting Reward Processing, Impulsivity, and Specific Aspects of Cocaine Addiction-Like Behavior in a Sex-Dependent Manner. Int J Neuropsychopharmacol, 2021. 24(11): p. 920–933.

8. Kaidanovich-Beilin, O., et al., Assessment of social interaction behaviors. J Vis Exp, 2011(48).

9. Dowell, D., et al., Prescribing Opioids for Pain — The New CDC Clinical Practice Guideline. New England Journal of Medicine, 2022. 387(22): p. 2011–2013.

10. Ferland, C.E., E. Vega, and P.M. Ingelmo, Acute pain management in children: challenges and recent improvements. Curr Opin Anaesthesiol, 2018. 31(3): p. 327–332.

11. Hudgins, J.D., et al., Trends in Opioid Prescribing for Adolescents and Young Adults in Ambulatory Care Settings. Pediatrics, 2019. 143(6).

12. Foster, A.A., et al., The use of opioids in low acuity pediatric trauma patients. PLoS One, 2019. 14(12): p. e0226433.

13. Rosoff, D.B., G.D. Smith, and F.W. Lohoff, Prescription Opioid Use and Risk for Major Depressive Disorder and Anxiety and Stress-Related Disorders: A Multivariable Mendelian Randomization Analysis. JAMA Psychiatry, 2021. 78(2): p. 151–160.

14. Scherrer, J.F., et al., Prescription Opioid Duration, Dose, and Increased Risk of Depression in 3 Large Patient Populations. Ann Fam Med, 2016. 14(1): p. 54–62.

15. Naji, L., et al., The association between age of onset of opioid use and comorbidity among opioid dependent patients receiving methadone maintenance therapy. Addiction Science & Clinical Practice, 2017. 12(1): p. 9.

16. Bell, T.M., et al., Outpatient Opioid Prescriptions are Associated With Future Substance Use Disorders and Overdose Following Adolescent Trauma. Ann Surg, 2022. 276(6): p. e955–e960.

17. Wilson, J.D., et al., Trajectories of Opioid Use Following First Opioid Prescription in Opioid-Naive Youths and Young Adults. JAMA Netw Open, 2021. 4(4): p. e214552.

18. Quinn, P.D., et al., Association of Opioid Prescription Initiation During Adolescence and Young Adulthood With Subsequent Substance-Related Morbidity. JAMA Pediatrics, 2020. 174(11): p. 1048–1055.

19. Miech, R., et al., Prescription Opioids in Adolescence and Future Opioid Misuse. Pediatrics, 2015. 136(5): p. e1169–77.

20. Gaw, C.E., et al., Characteristics of Fatal Poisonings Among Infants and Young Children in the United States. Pediatrics, 2023.

21. Yule, A.M., R.M. Lyons, and T.E. Wilens, Opioid Use Disorders in Adolescents-Updates in Assessment and Management. Curr Pediatr Rep, 2018. 6(2): p. 99–106.

22. Lamblin, M., et al., Social connectedness, mental health and the adolescent brain. Neurosci Biobehav Rev, 2017. 80: p. 57–68.

23. Sakata, J.T., I. Catalano, and S.C. Woolley, Mechanisms, development, and comparative perspectives on experience-dependent plasticity in social behavior. J Exp Zool A Ecol Integr Physiol, 2022. 337(1): p. 35–49.

24. Vanderschuren, L.J. and V. Trezza, What the laboratory rat has taught us about social play behavior: role in behavioral development and neural mechanisms. Curr Top Behav Neurosci, 2014. 16: p. 189–212.

25. Andrews, J.L., S.P. Ahmed, and S.J. Blakemore, Navigating the Social Environment in Adolescence: The Role of Social Brain Development. Biol Psychiatry, 2021. 89(2): p. 109–118.

26. Kopec, A.M., et al., Microglial dopamine receptor elimination defines sex-specific nucleus accumbens development and social behavior in adolescent rats. Nat Commun, 2018. 9(1): p. 3769.

27. Chu, X., et al., Sexual incentive motivation and male and female copulatory behavior in female rats given androgen from postnatal day 20. Physiol Behav, 2021. 237: p. 113460.

28. Hunt, T.K.A., K.S. Slack, and L.M. Berger, Adverse childhood experiences and behavioral problems in middle childhood. Child Abuse Negl, 2017. 67: p. 391–402.

29. Pierce, H., M.S. Jones, and E.A. Holcombe, Early Adverse Childhood Experiences and Social Skills Among Youth in Fragile Families. J Youth Adolesc, 2022. 51(8): p. 1497–1510.

30. Potrebić, M., et al., The Influence of Social Isolation on Social Orientation, Sociability, Social Novelty Preference, and Hippocampal Parvalbumin-Expressing Interneurons in Peripubertal Rats - Understanding the Importance of Meeting Social Needs in Adolescence. Front Behav Neurosci, 2022. 16: p. 872628.

31. Gartland, L.A., et al., Sociability as a personality trait in animals: methods, causes and consequences. Biol Rev Camb Philos Soc, 2022. 97(2): p. 802–816.

32. Boswell, N., et al., A review and preview of developments in the measurement of sociability. Bull Menninger Clin, 2020. 84(1): p. 79–101.

33. Caldwell, H.K., Neurobiology of sociability. Adv Exp Med Biol, 2012. 739: p. 187–205.

34. Lopez-Tobon, A., S. Trattaro, and G. Testa, The sociability spectrum: evidence from reciprocal genetic copy number variations. Mol Autism, 2020. 11(1): p. 50.

35. Yizhar, O. and D.R. Levy, The social dilemma: prefrontal control of mammalian sociability. Curr Opin Neurobiol, 2021. 68: p. 67–75.

36. Bales, K.L., et al., Social touch during development: Long-term effects on brain and behavior. Neurosci Biobehav Rev, 2018. 95: p. 202–219.

37. Saarinen, A., et al., Social touch experience in different contexts: A review. Neurosci Biobehav Rev, 2021. 131: p. 360–372.

38. Wu, Y.E., et al., Neural control of affiliative touch in prosocial interaction. Nature, 2021. 599(7884): p. 262–267.

39. Jablonski, N.G., Social and affective touch in primates and its role in the evolution of social cohesion. Neuroscience, 2021. 464: p. 117–125.

40. Hunnius, S., The early development of visual attention and its implications for social and cognitive development. Prog Brain Res, 2007. 164: p. 187–209.

41. Tsang, T., et al., Social complexity and the early social environment affect visual social attention to faces. Autism Res, 2019. 12(3): p. 445–457.

42. Dindar, K., et al., Social-Pragmatic Inferencing, Visual Social Attention and Physiological Reactivity to Complex Social Scenes in Autistic Young Adults. J Autism Dev Disord, 2022. 52(1): p. 73–88.

43. Wang, L. and R.J. Krauzlis, Visual Selective Attention in Mice. Curr Biol, 2018. 28(5): p. 676–685.e4.

44. Meier, I.M., et al., A mu-opioid feedback model of human social behavior. Neuroscience & Biobehavioral Reviews, 2021. 121: p. 250–258.

45. Lee, M., et al., A comprehensive review of opioid-induced hyperalgesia. Pain Physician, 2011. 14(2): p. 145–61.

46. Kadhim, S., J. McDonald, and D.G. Lambert, Opioids, gliosis and central immunomodulation. J Anesth, 2018. 32(5): p. 756–767.

47. Gardner, E.L., Addiction and brain reward and antireward pathways. Adv Psychosom Med, 2011. 30: p. 22–60.

48. King’uyu, D.N., S.B.Z. Stephens, and A.M. Kopec, Immune signaling in sex-specific neural and behavioral development: Adolescent opportunity. Curr Opin Neurobiol, 2022. 77: p. 102647.

49. Peper, J.S., S.M. Burke, and L.M. Wierenga, Sex differences and brain development during puberty and adolescence. Handb Clin Neurol, 2020. 175: p. 25–54.

50. Butler, G. and P. Purushothaman, Delayed puberty. Minerva Pediatr, 2020. 72(6): p. 484–490.

51. Mendle, J., et al., Understanding Puberty and Its Measurement: Ideas for Research in a New Generation. J Res Adolesc, 2019. 29(1): p. 82–95.

52. Vijayakumar, N., et al., Puberty and the human brain: Insights into adolescent development. Neurosci Biobehav Rev, 2018. 92: p. 417–436.

53. Euling, S.Y., et al., Examination of US puberty-timing data from 1940 to 1994 for secular trends: panel findings. Pediatrics, 2008. 121 Suppl 3: p. S172–91.

54. Reddy, R.B., A.A. Sandel, and R.E. Dahl, Puberty initiates a unique stage of social learning and development prior to adulthood: Insights from studies of adolescence in wild chimpanzees. Dev Cogn Neurosci, 2022. 58: p. 101176.

55. Pfeifer, J.H. and N.B. Allen, Puberty Initiates Cascading Relationships Between Neurodevelopmental, Social, and Internalizing Processes Across Adolescence. Biol Psychiatry, 2021. 89(2): p. 99–108.

56. Stuart, S.A. and E.S. Robinson, Reducing the stress of drug administration: implications for the 3Rs. Sci Rep, 2015. 5: p. 14288.

57. Turner, P.V., et al., Administration of substances to laboratory animals: routes of administration and factors to consider. J Am Assoc Lab Anim Sci, 2011. 50(5): p. 600–13.

58. Casey, B.J., et al., The Adolescent Brain Cognitive Development (ABCD) study: Imaging acquisition across 21 sites. Dev Cogn Neurosci, 2018. 32: p. 43–54.

59. Dick, A.S., et al., Meaningful associations in the adolescent brain cognitive development study. Neuroimage, 2021. 239: p. 118262.

60. Hagler, D.J., Jr., et al., Image processing and analysis methods for the Adolescent Brain Cognitive Development Study. Neuroimage, 2019. 202: p. 116091.

61. Dayananda, K.K., et al., Early life stress impairs synaptic pruning in the developing hippocampus. Brain Behav Immun, 2023. 107: p. 16–31.

62. Evans, G.W., D. Li, and S.S. Whipple, Cumulative risk and child development. Psychol Bull, 2013. 139(6): p. 1342–96.

63. McLaughlin, K.A. and M.A. Sheridan, Beyond Cumulative Risk: A Dimensional Approach to Childhood Adversity. Curr Dir Psychol Sci, 2016. 25(4): p. 239–245.

64. McLaughlin, K.A., et al., Mechanisms linking childhood trauma exposure and psychopathology: a transdiagnostic model of risk and resilience. BMC Med, 2020. 18(1): p. 96.

65. Stephens, S.B. and K. Wallen, Environmental and social influences on neuroendocrine puberty and behavior in macaques and other nonhuman primates. Horm Behav, 2013. 64(2): p. 226–39.

66. Horwitz, A.V., Social Context, Biology, and the Definition of Disorder. J Health Soc Behav, 2017. 58(2): p. 131–145.

67. Bredewold, R., et al., Sex-specific modulation of juvenile social play behavior by vasopressin and oxytocin depends on social context. Front Behav Neurosci, 2014. 8: p. 216.

68. Veenema, A.H., R. Bredewold, and G.J. De Vries, Sex-specific modulation of juvenile social play by vasopressin. Psychoneuroendocrinology, 2013. 38(11): p. 2554–2561.

69. Thomas, P.A., H. Liu, and D. Umberson, Family Relationships and Well-Being. Innov Aging, 2017. 1(3): p. igx025.

70. Toyoshima, A. and J. Nakahara, The Effects of Familial Social Support Relationships on Identity Meaning in Older Adults: A Longitudinal Investigation. Front Psychol, 2021. 12: p. 650051.

71. Daley, D.C., Family and social aspects of substance use disorders and treatment. J Food Drug Anal, 2013. 21(4): p. S73–s76.

72. Krause, M.A. and R.W. Mitchell, Object-Choice Test, in Encyclopedia of Animal Cognition and Behavior, J. Vonk and T. Shackelford, Editors. 2018, Springer International Publishing: Cham. p. 1–12.

73. Kraeuter, A.K., P.C. Guest, and Z. Sarnyai, Free Dyadic Social Interaction Test in Mice. Methods Mol Biol, 2019. 1916: p. 93–97.

74. Schiavi, S., et al., Assessing Dyadic Social Interactions in Rodent Models of Neurodevelopmental Disorders, in Translational Research Methods in Neurodevelopmental Disorders, S. Martin and F. Laumonnier, Editors. 2022, Springer US: New York, NY. p. 193–216.

75. Morozov, A., Behavioral Modulation by Social Experiences in Rodent Models. Curr Protoc Neurosci, 2018. 84(1): p. e50.

76. Kiyokawa, Y., et al., Social buffering reduces male rats’ behavioral and corticosterone responses to a conditioned stimulus. Horm Behav, 2014. 65(2): p. 114–8.

77. Denommé, M.R. and G.J. Mason, Social Buffering as a Tool for Improving Rodent Welfare. J Am Assoc Lab Anim Sci, 2022. 61(1): p. 5–14.

78. Perich, T., P.B. Mitchell, and B. Vilus, Stigma in bipolar disorder: A current review of the literature. Aust N Z J Psychiatry, 2022. 56(9): p. 1060–1064.

79. Zweifel, P., Mental health: The burden of social stigma. Int J Health Plann Manage, 2021. 36(3): p. 813–825.

80. Sasson, N.J., et al., Neurotypical Peers are Less Willing to Interact with Those with Autism based on Thin Slice Judgments. Sci Rep, 2017. 7: p. 40700.

81. Derrick, J.L., L.D. Wittkower, and J.D. Pierce, Committed relationships and substance use: recent findings and future directions. Curr Opin Psychol, 2019. 30: p. 74–79.

82. Cadigan, J.M., et al., Influence of developmental social role transitions on young adult substance use. Curr Opin Psychol, 2019. 30: p. 87–91.

83. Sterrett-Hong, E.M., et al., The Impact of Closeness to Non-Parental Adults in Social Networks on Substance Use among Young Men Who Have Sex with Men. J Homosex, 2021. 68(10): p. 1727–1744.

84. Ruthig, P. and M. Schönwiesner, Common principles in the lateralization of auditory cortex structure and function for vocal communication in primates and rodents. Eur J Neurosci, 2022. 55(3): p. 827–845.

85. Ebbesen, C.L. and R.C. Froemke, Body language signals for rodent social communication. Curr Opin Neurobiol, 2021. 68: p. 91–106.

86. Rymer, T.L., The Role of Olfactory Genes in the Expression of Rodent Paternal Care Behavior. Genes (Basel), 2020. 11(3).

87. Venniro, M. and Y. Shaham, An operant social self-administration and choice model in rats. Nat Protoc, 2020. 15(4): p. 1542–1559.

88. Venniro, M., et al., Volitional social interaction prevents drug addiction in rat models. Nat Neurosci, 2018. 21(11): p. 1520–1529.

89. Venniro, M., et al., Operant Social Reward Decreases Incubation of Heroin Craving in Male and Female Rats. Biol Psychiatry, 2019. 86(11): p. 848–856.

90. McCrory, E.J. and L. Mayes, Understanding Addiction as a Developmental Disorder: An Argument for a Developmentally Informed Multilevel Approach. Curr Addict Rep, 2015. 2(4): p. 326–330.

91. Compton, W.M., E.M. Wargo, and N.D. Volkow, Neuropsychiatric Model of Addiction Simplified. Psychiatr Clin North Am, 2022. 45(3): p. 321–334.

92. Strashny, A., Age of Substance Use Initiation Among Treatment Admissions Aged 18 to 30, in The CBHSQ Report. 2013, Substance Abuse and Mental Health Services Administration (US): Rockville (MD). p. 1–9.

93. Chen, C.Y., C.L. Storr, and J.C. Anthony, Early-onset drug use and risk for drug dependence problems. Addict Behav, 2009. 34(3): p. 319–22.

94. Mantey, D.S., et al., The association between age of initiation and current blunt use among adults: Findings from the national survey on drug use and health, 2014-2019. Drug Alcohol Depend, 2022. 230: p. 109191.

95. Giletta, M., et al., A meta-analysis of longitudinal peer influence effects in childhood and adolescence. Psychol Bull, 2021. 147(7): p. 719–747.

96. Barbu, S., G. Cabanes, and G. Le Maner-Idrissi, Boys and girls on the playground: sex differences in social development are not stable across early childhood. PLoS One, 2011. 6(1): p. e16407.

97. Kyle, S.C., G.M. Burghardt, and M.A. Cooper, Development of social play in hamsters: Sex differences and their possible functions. Brain Res, 2019. 1712: p. 217–223.

98. Strickland, J.C. and M.A. Smith, The effects of social contact on drug use: behavioral mechanisms controlling drug intake. Exp Clin Psychopharmacol, 2014. 22(1): p. 23–34.

99. El Rawas, R., I.M. Amaral, and A. Hofer, Social interaction reward: A resilience approach to overcome vulnerability to drugs of abuse. European Neuropsychopharmacology, 2020. 37: p. 12–28.

100. Chen, G., Social support, spiritual program, and addiction recovery. Int J Offender Ther Comp Criminol, 2006. 50(3): p. 306–23.

101. Lookatch, S.J., A.S. Wimberly, and J.R. McKay, Effects of Social Support and 12-Step Involvement on Recovery among People in Continuing Care for Cocaine Dependence. Subst Use Misuse, 2019. 54(13): p. 2144–2155.

102. Redhead, D. and E.A. Power, Social hierarchies and social networks in humans. Philos Trans R Soc Lond B Biol Sci, 2022. 377(1845): p. 20200440.

103. Chen, S., Social power and the self. Curr Opin Psychol, 2020. 33: p. 69–73.

104. Kim, B., et al., Endocrine disruptors alter social behaviors and indirectly influence social hierarchies via changes in body weight. Environ Health, 2015. 14: p. 64.

105. Frank, D., et al., Assessing Dominant-Submissive Behavior in Adult Rats Following Traumatic Brain Injury. J Vis Exp, 2022(190).

106. Malatynska, E. and R.J. Knapp, Dominant-submissive behavior as models of mania and depression. Neurosci Biobehav Rev, 2005. 29(4-5): p. 715–37.

107. Wise, R.A. and M.A. Robble, Dopamine and Addiction. Annu Rev Psychol, 2020. 71: p. 79–106.

108. Pomrenze, M.B., F. Paliarin, and R. Maiya, Friend of the Devil: Negative Social Influences Driving Substance Use Disorders. Front Behav Neurosci, 2022. 16: p. 836996.

109. Pomrenze, M.B., et al., Modulation of 5-HT release by dynorphin mediates social deficits during opioid withdrawal. Neuron, 2022. 110(24): p. 4125–4143.e6.

110. Delfs, J.M., et al., Noradrenaline in the ventral forebrain is critical for opiate withdrawal-induced aversion. Nature, 2000. 403(6768): p. 430–4.

111. Piccin, A. and A. Contarino, Long-lasting pseudo-social aggressive behavior in opiate-withdrawn mice. Progress in Neuro-Psychopharmacology and Biological Psychiatry, 2020. 97: p. 109780.

112. Stephan, A.H., B.A. Barres, and B. Stevens, The complement system: an unexpected role in synaptic pruning during development and disease. Annu Rev Neurosci, 2012. 35: p. 369–89.

113. Schafer, D.P., et al., Microglia sculpt postnatal neural circuits in an activity and complement-dependent manner. Neuron, 2012. 74(4): p. 691–705.

114. Stevens, B., et al., The classical complement cascade mediates CNS synapse elimination. Cell, 2007. 131(6): p. 1164–78.

115. Evrard, M.R., et al., Preventing adolescent synaptic pruning in mouse prelimbic cortex via local knockdown of α4βδ GABA(A) receptors increases anxiety response in adulthood. Sci Rep, 2021. 11(1): p. 21059.

116. Paolicelli, R.C., et al., Synaptic pruning by microglia is necessary for normal brain development. Science, 2011. 333(6048): p. 1456–8.

117. King’uyu, D.N., et al., The effect of morphine on rat microglial phagocytic activity: an in vitro study of brain region-, plating density-, sex-, morphine concentration-, and receptor-dependency. bioRxiv, 2023: p. 2022.10.03.510683.

118. Fredriksson, I., et al., Animal Models of Drug Relapse and Craving after Voluntary Abstinence: A Review. Pharmacol Rev, 2021. 73(3): p. 1050–1083.

119. Spanagel, R., Animal models of addiction. Dialogues Clin Neurosci, 2017. 19(3): p. 247–258.

120. Hyman, S.E., R.C. Malenka, and E.J. Nestler, Neural mechanisms of addiction: the role of reward-related learning and memory. Annu Rev Neurosci, 2006. 29: p. 565–98.

121. Jones, K., et al., Opioid Reduction Through Postoperative Pain Management in Pediatric Orthopedic Surgery. Pediatrics, 2021. 148(6).

122. Gaglani, A. and T. Gross, Pediatric Pain Management. Emerg Med Clin North Am, 2018. 36(2): p. 323–334.

123. Báez-Mendoza, R., et al., Neuronal Circuits for Social Decision-Making and Their Clinical Implications. Front Neurosci, 2021. 15: p. 720294.

124. Terenzi, D., et al., Determinants and modulators of human social decisions. Neuroscience & Biobehavioral Reviews, 2021. 128: p. 383–393.

125. Wegmann, E., et al., Social-networks-related stimuli interferes decision making under ambiguity: Interactions with cue-induced craving and problematic social-networks use. J Behav Addict, 2021. 10(2): p. 291–301.

126. Volkow, N.D., et al., Unbalanced neuronal circuits in addiction. Curr Opin Neurobiol, 2013. 23(4): p. 639–48.

127. Volkow, Nora D., Ruben D. Baler, and Rita Z. Goldstein, Addiction: Pulling at the Neural Threads of Social Behaviors. Neuron, 2011. 69(4): p. 599–602.

